# Association versus Prediction: the impact of cortical surface smoothing and parcellation on brain age

**DOI:** 10.1101/2020.11.29.403105

**Authors:** Yashar Zeighami, Alan C. Evans

**Affiliations:** Montreal Neurological Institute, McGill University, Montreal, Quebec, Canada; Ludmer Centre for Neuroinformatics and Mental Health, McGill University, Montreal, Quebec, Canada

**Keywords:** Brain aging, Cortical Thickness, Prediction, Delta age, Smoothing, Parcellation

## Abstract

Association and prediction studies of the brain target the biological consequences of aging and their impact on brain function. Such studies are conducted using different smoothing levels and parcellations at the preprocessing stage, on which their results are dependent. However, the impact of these parameters on the relationship between association values and prediction accuracy is not established. In this study, we used cortical thickness and its relationship with age to investigate how different smoothing and parcellation levels affect the detection of age-related brain correlates as well as brain age prediction accuracy. Our main measures were resel numbers - resolution elements - and age-related variance explained. Using these common measures enabled us to directly compare parcellation and smoothing effects in both association and prediction studies. In our sample of N=608 participants with age range 18-88, we evaluated age-related cortical thickness changes as well as brain age prediction. We found a negative relationship between prediction performance and correlation values for both parameters. Our results also quantify the relationship between delta age estimates obtained based on different processing parameters. Furthermore, with the direct comparison of the two approaches, we highlight the importance of correct choice of smoothing and parcellation parameters in each task, and how they can affect the results of the analysis in opposite directions.

## 1 Introduction

From a biological standpoint, aging is defined by the structural and functional alterations in living organisms (López-Otín et al., 2013). Traditionally, brain imaging studies have used neuroimaging data to find associations between age and tissue alterations across brain areas, using chronological age as the ground truth (Booth et al., 2013; Curiati et al., 2009; Hu et al., 2014; Lemaître et al., 2005; Takahashi et al., 2011; Ziegler et al., 2012). However, biological age might vary between individuals with identical chronological age as well as across different tissues within the same person (Horvath, 2013). To non-invasively measure the biological age of the brain, neuroimaging data is used to predict age. The difference between predicted age and chronological age is then defined as “delta” or brain age gap estimate i.e. “BrainAGE” to compare the subjects’ chronological age with the predicted brain age in a given reference population (James H. Cole & Franke, 2017; Franke et al., 2012; Franke & Gaser, 2019a; Smith et al., 2019a).

Both age related brain alterations and delta age have been studied and used extensively in the neuroimaging literature. Age association studies translate and generalize easily across different datasets. These association studies are applied across brain regions and can distinguish the differential effect of age on different brain areas (Storsve et al., 2014). Furthermore, they directly relate to biological measures and mechanistic changes in the brain (Khundrakpam et al., 2015). More recently, it has been recognized that association studies are prone to overfitting and more studies focus on prediction as the main goal of the study (Bzdok et al., 2020; Yarkoni & Westfall, 2017). Brain age studies (i.e. age prediction studies based on neuroimaging data) rely on modeling and prediction accuracy. This goal is generally achieved by using a feature set that can capture the variability between and within subjects. On the other hand, prediction tasks face a trade-off between a more accurate whole brain model with no regional specificity versus a model with lower accuracy and increased spatial resolution (James H. Cole & Franke, 2017; Franke & Gaser, 2019a). This limitation also results in a more indirect relationship between delta age and other phenotypes without a direct mechanistic and biological model. Nonetheless, the difference between brain age and chronological age is associated with cognitive decline (Gaser et al., 2013), predisposition to neuropsychiatric and neurodegenerative disorders (Kaufmann et al., 2019), and mortality (J. H. Cole et al., 2018). While evidence supports the application of delta age as a valuable measure to study aging in health and disease, it has been criticized due to its reliance on prediction accuracy (i.e. more accurate models result in lower delta values) (James H. Cole & Franke, 2017).

The results of both association studies and delta estimation studies are impacted by processing steps such as data normalization, spatial resolution, and parcellation level (i.e. size of the parcels) of the analysis. Most association studies use smoothing to (i) normalize the distributions of cortical thickness across subjects, (ii) minimize registration and anatomical misalignment across subjects, (iii) reduce measurement noise, and (iv) increase statistical power (Lerch et al., 2006; Lerch & Evans, 2005; Worsley et al., 1999; Zhao et al., 2013). These advantages are gained at the cost of losing individual variability and spatial resolution. In fact, smoothing has been studied and optimized for best performance in association studies, using simulation as well as in real datasets. The smoothing level has been proposed as a dimension within the parameter space in the association analysis that needs to be searched for the given statistical contrast (Lerch & Evans, 2005; Zhao et al., 2013).

Brain age prediction studies have been conducted with various levels of data smoothing. Moreover, these studies rely on various dimension reduction techniques, brain parcellations, or a combination of the two approaches for feature extraction (Franke & Gaser, 2019b; Smith et al., 2019b). The optimal parcellation for a given task is an open research topic and it can vary between studies (Eickhoff et al., 2018; Gorgolewski et al., 2016; Salehi et al., 2020). While some studies have predicted brain age with multiple parcellation resolutions (Khundrakpam et al., 2015; J. D. Lewis et al., 2019), others have used a predetermined number of parcels. However, the effect of smoothing and parcellation in brain age prediction is not studied systematically. Furthermore, these changes in prediction accuracy also affect the delta estimate (i.e. the variable of interest), and it is not clear whether the delta estimates are robust or sensitive toward these initial choices.

In this study, we used cortical thickness as the brain measure of interest and examined the effect of smoothing and parcellation level on both brain associations with age and brain age prediction. Using different levels of parcellation and smoothing, we projected brain measures onto a lower dimension data representation space and investigated how this mapping affects the derived associations and predictions. We further examined the relationship between the two approaches. Finally, we examined how delta age estimates alter based on different smoothing and parcellation levels.

## 2 Methods

### 2.1 Data

Data used in this study included subjects with T1-weighted MRI data available from the second stage of the Cambridge Centre for Ageing and Neuroscience (CamCAN, https://www.cam-can.org/index.php?content=dataset) dataset, described in more detail in (Shafto et al., 2014; Taylor et al., 2017). Subjects were screened for neurological and psychiatric conditions and those with such underlying disorders were excluded from the study.

### 2.2 MRI acquisition

T1-weighted MRIs were acquired on a 3T Siemens TIM Trio, with a 32 channel head-coil using a 3D magnetization-prepared rapid gradient echo (MPRAGE) sequence (TR=2250ms, TE=2.99ms, TI=900ms; FA=9 deg; FOV=256×240×192mm; 1mm isotropic; GRAPPA=2; TA=4mins 32s). For detailed acquisition parameters see: https://camcan-archive.mrc-cbu.cam.ac.uk/dataaccess/pdfs/CAMCAN700_MR_params.pdf.

### 2.3 MRI processing

We used CIVET 2.1.1 (http://www.bic.mni.mcgill.ca/ServicesSoftware/CIVET, release December 2019), a fully automated structural image analysis pipeline developed at the Montreal Neurological Institute, to perform surface extraction and cortical thickness estimation. Briefly, each subject’s T1-weighted MRI is corrected for nonuniformity artifacts using the N3 algorithm (N3 distance = 125 mm) (Sled et al., 1998) and linearly registered to stereotaxic MNI152 space (voxel resolution = 0.5 mm) (Collins et al., 1994). The brain is extracted and undergoes tissue classification into three classes: white matter (WM) tissue, grey matter (GM) tissue, and cerebrospinal fluid (CSF) (Tohka et al., 2004; Zijdenbos et al., 2002). White and grey matter surfaces are extracted using the marching cube algorithm and constrained Laplacian-based automated segmentation with proximities (CLASP) algorithms, respectively (Kabani et al., 2001; Kim et al., 2005; MacDonald et al., 2000). Using the extracted surfaces, cortical thickness is measured as the distance between the white and grey cortical surfaces using the Laplace’s equation (Jones et al., 2000). For blurring, a surface-based diffusion smoothing kernel (not to be confused with volumetric kernels) is used, which generalizes Gaussian kernel smoothing and applies it to the curved cortical surfaces (Chung et al., 2002). We applied 6 different smoothing levels with FWHM = 0, 5, 10, 20, 30, and 40 mm. Cortical thickness was measured across the cortical surface for 81924 vertices (40962 vertices per hemisphere). The results underwent visual inspection, specifically subjects with major errors in extracted pial and gray–white surfaces were excluded.

### 2.4 Cortical parcellations

We used the Schaefer functional MRI parcellations (Schaefer et al., 2018), a data-driven atlas based on the widely used seven large-scale functional network parcellations by (Thomas Yeo et al., 2011). We used Schaefer parcellation with 100, 200, 400, and 1000 regions (referred to as parcellation levels). All atlases were registered to the MNI cortical surface template and used in the MNI space (L. B. Lewis et al., 2019). Cortical thickness measurements with different smoothing levels were averaged across these parcellations. These parcellation based measures of cortical thickness were used alongside vertex-wise measurements to examine the interaction between the effect of brain parcellation averaging and smoothing on statistical associations as well as brain age prediction accuracies.

### 2.5 Cortical resels and effective smoothing

In order to compare the findings between smoothing levels and different parcellations, first all obtained cortical thickness were projected to the brain surface. We used the number of resels (i.e. resolution elements) as the measure of interest, since it takes the statistical dependence of the brain map into consideration and is independent of the analysis resolution (at least from a theoretical standpoint) (Lerch et al., 2006; Worsley, 1996; Worsley et al., 1992, 1999). Using the statistical maps between aging and cortical thickness, we estimated the number of resels for each smoothing and parcellation level and used it to quantify the similarity between these conditions. Resels are the number of resolution elements approximated for a given search space (i.e. D(S2), S2= brain surface) and a given smoothness level FWHM. While the effective FWHM measure varies across brain areas, we defined the overall effective smoothness of the brain map as the square root of the surface search space divided by the number of resels estimated across brain areas (Hayasaka et al., 2004). For the purpose of the current study, the main statistical maps considered are the linear associations between cortical thickness and the chronological age of the participants. All analysis were performed using SurfStat toolbox https://www.math.mcgill.ca/keith/surfstat/.

### 2.6 Statistical methods

To examine the effect of the smoothing and parcellations, mean (μ) and standard deviation (σ) of cortical thickness for each vertex/parcel was calculated across the population. The coefficient of variation (CV), *CV* = σ/μ, was used as the main measure of variability. The CV was averaged across the 7 main cytoarchitectural brain regions (von Economo, CF; Koskinas, 1927) in order to examine the effect of parcellation and smoothing across major cytoarchitectural regions and identify any differential impact on a given brain region. Finally, to measure the association between chronological age and cortical thickness across lifespan, correlation coefficient (*r*) for each vertex/region was calculated. Variance explained (*r*^2^) was used to visualize the results.

### 2.7 Brain age prediction

We used principal component analysis (PCA), a singular value decomposition based data factorization method, as the dimensionality reduction approach for our predictive variables (i.e. cortical thickness data) (Smith et al., 2019b). This approach allowed us to use the same number of features across parcellation levels and smoothing kernels and therefore made it possible to compare model performance across these conditions. Our analysis for each condition included 1 to 100 first principal components as features to study different levels of dimensionality reduction. 100 is used as the maximum possible number of independent components for the lowest number of parcels (i.e. Schaefer 100). To predict brain age, we used linear regression as the main prediction model, and to ensure generalizability and avoid overfitting, we used 10-fold cross validation. Finally, to increase robustness, results averaged over 100 repetitions are reported. Root-mean-squared error (RMSE) was used as the natural cost function for linear regression models. Mean absolute error (MAE) and correlation between chronological age and predicted age (two other common error metrics in the age prediction literature (Franke & Gaser, 2019b)) are also reported in the supplementary materials.

### 2.8 The relationship between Brain age prediction and age related brain association

To compare brain age association and age prediction, we used the variance explained between dependent and independent variables as the main measure of interest for each model. This common measure enabled us to quantify the two analyses in relation to each other. Furthermore, we examined how the number of resels affects whole brain associations with age as well as brain age prediction. To translate the age prediction error into variance explained, we used the predictive features in a linear model, calculating the variance explained for age using adjusted *R*^2^. Finally, the overfitting bias between the variance explained (i.e. adjusted *R*^2^) using this linear model and the cross validated prediction (i.e. *r*^2^ between predicted age and chronological age) is reported.

### 2.9 Delta age

The main goal of brain age prediction studies is to calculate the deviation from chronological age based on the population norm, also known as delta age. Here, we examined the effect of smoothing and parcellation on delta age estimation:

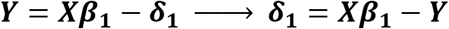

where Y denotes chronological age, X denotes the neuroimaging features, and **δ_1_** denotes the difference between predicted and chronological age. **δ_1_** is a measure of brain state/health compared to the population with similar chronological age, and is used to study the predisposition to different brain disorders as well as individual cognitive abilities in neuroimaging literature.

**δ_1_** being residual of the predictive model is by definition: (1) orthogonal to the predictive measures *X*, and in the case of linear models (2) correlated with the output *Y*(i.e. chronological age) (Le et al., 2018; Liang et al., 2019; Smith et al., 2019b). The first feature is unfavorable, since we are interested in brain related discrepancy between chronological and predicted age. The lack of association between **δ_1_** and brain features predicting age undermines the interpretability of **δ_1_** in relation to brain measures. The second property is also an adverse feature, since it makes it difficult to distinguish the effect of the chronological age from the additional biological delta age (due to their collinearity). Therefore, in the current study, we followed the recommendation of smith and colleagues (Smith et al., 2019b) and used **δ_2_**, the orthogonalized residuals against chronological age:

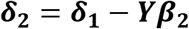

**δ_2_** is then used as the main measure of interest for association across conditions. The results for **δ_1_** is provided in the supplementary materials. Note that **δ_2_** is also consistently calculated using the same 10-fold cross validation with 100 repeats as **δ_1_**. All statistical and prediction analyses were performed using MATLAB 2018a.

## 3 Results

### 3.1 Cortical thickness aging, resels and practical smoothness

The parcellations have a considerable impact on the number of resels and function as region-based smoothing kernels applied across the brain (Figure 1-A). This change in the number of resels affects the statistical power and the association as well as prediction results. Across parcellation levels from 100 to 1000, the effect of the smaller smoothing kernels with FWHM 0-10 mm is negligible, while applying larger kernels reduces the number of resels dramatically. This equivalency plot also suggests that at the vertex level, the smoothing kernels act as a non-specific parcellation (from an anatomical perspective) across the brain.

**Figure 1.**
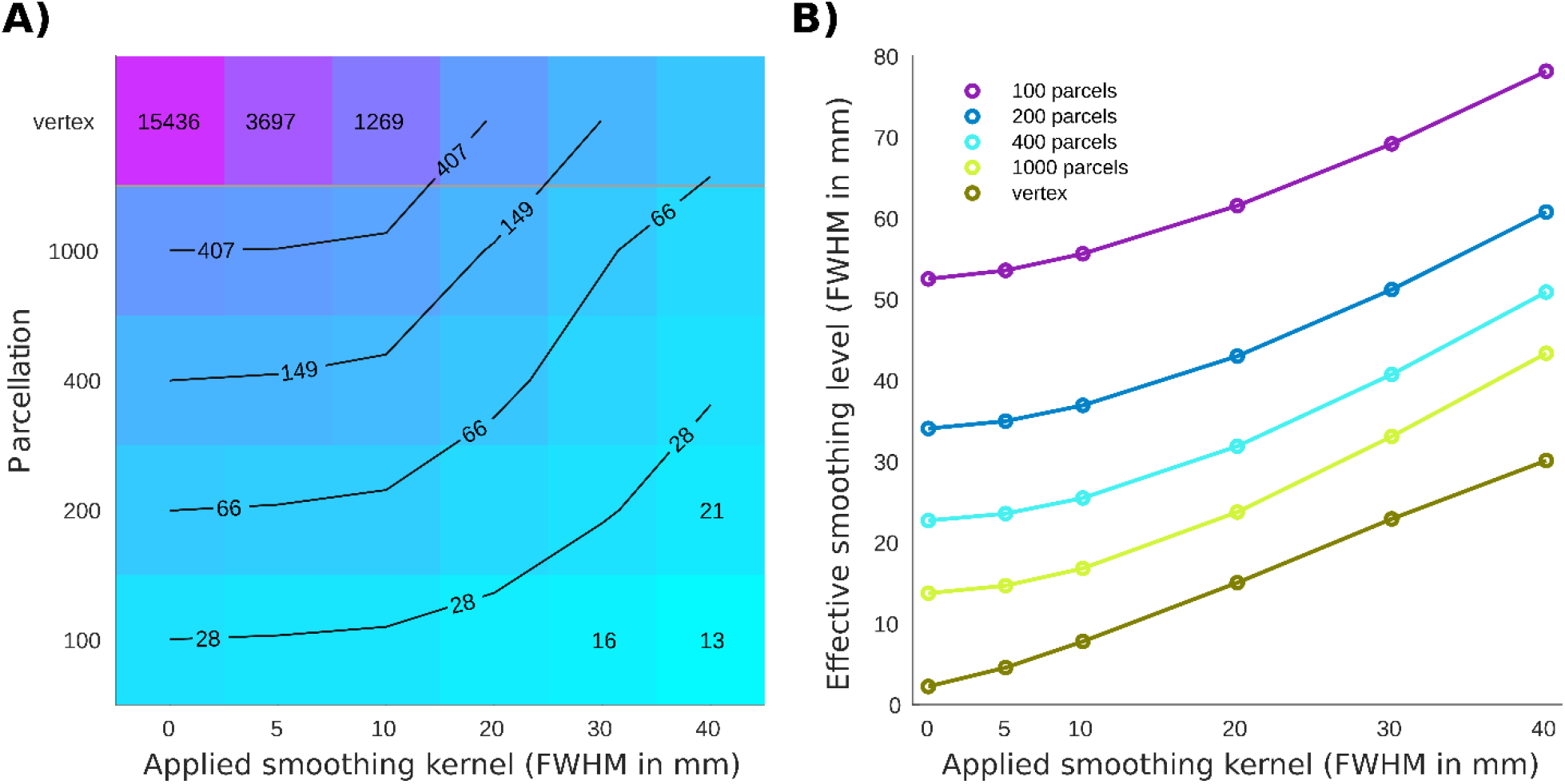
Number of resels and affective smoothing for cortical thickness association with age. **A)** Number of resels estimated for different parcellations/smoothing pairs. The lines show the interpolated iso-response values. **B)** Effective smoothing based on the number of resels for each condition. The results show the initial effective smoothing as a result of parcellation with additional smoothing with applied smoothing kernels.

### 3.2 Cortical thickness variability

While keeping the mean cortical thickness measure intact, smoothing resulted in underestimation of the cortical thickness in the gyri areas and overestimation in the sulci regions. The results are similar for parcellations in the case of uniformly sized parcels and balanced inclusion of gyri and sulci in each parcel (both criteria are met in Schaefer parcellations). Cortical thickness variability (i.e. CV) reduces significantly both as a result of using greater smoothing and larger parcels (Figure 2-A).

**Figure 2.**
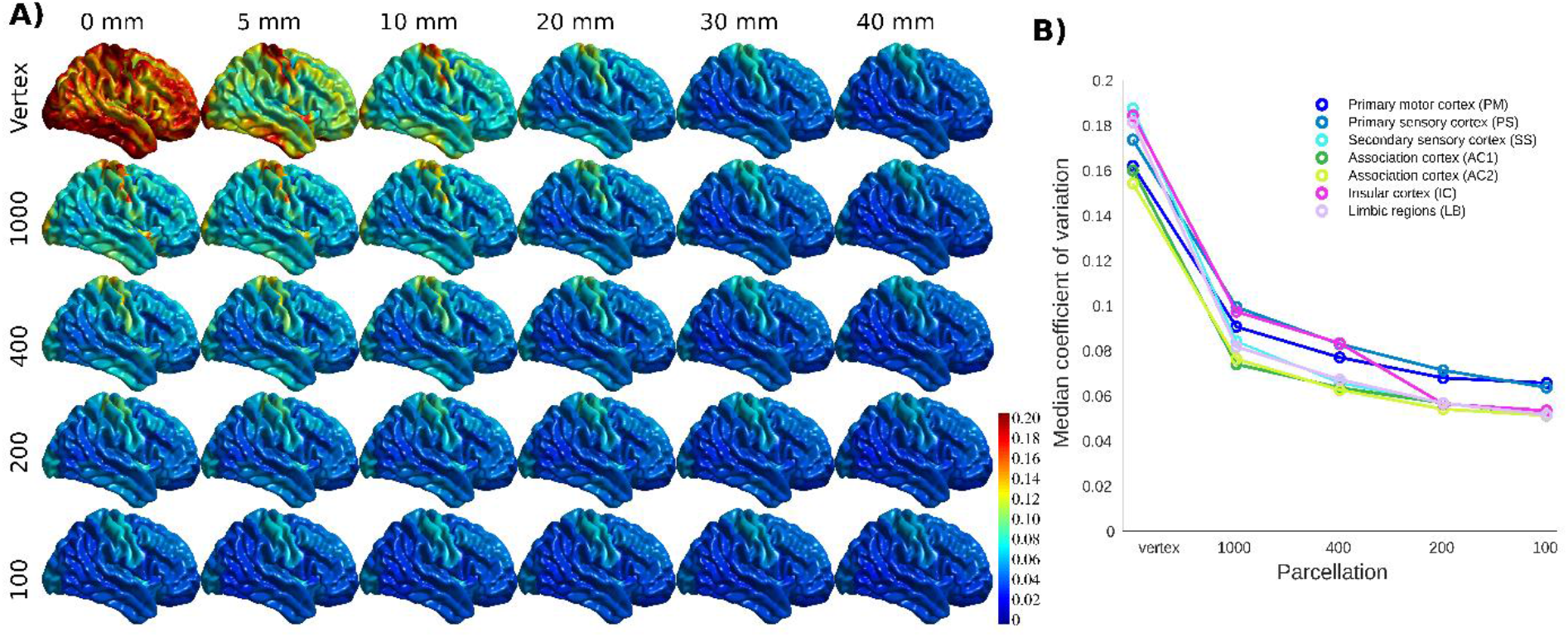
The coefficient of variation (CV) of cortical thickness across population. **A)** CV projected across brain vertices for each parcellations/smoothing pair. **B)** CV shown at 0 mm smoothing level for each cytoarchitectural region across parcellation resolutions.

The association cortices have the lowest CV across resolutions and parcellations. Both smoothing and parcellation result in the highest decrease in CV in limbic and insular cortices, while primary sensory and motor areas show the lowest change (Figure 2-B). The results are shown for 0 mm smoothing across parcellations. The greatest change occurs with increasing the FWHM value from 10 to 20 mm, as well as decreasing the number of parcels from 400 to 200. The results for different smoothing kernels at vertex level were also similar (Supplementary Figure 1).

### 3.3 statistical association between cortical thickness and aging

Figure 3-A. shows the association between age and cortical thickness (using variance explained *r*^2^), calculated for each voxel/parcel for all conditions, after Bonferroni correction to account for the multiple comparisons at each level. The correlation increases with greater smoothing and larger parcels. Changing smoothing kernel size results in the highest variability in the correlation distribution across the brain at vertex level resolution (Figure 3-B, top panel), whereas smoothing doesn’t change the results within Schaefer 100 parcellations (Figure 3-B, bottom panel). The same pattern is evident between parcellation levels with 0 mm smoothing showing the highest variability, and 40 mm smoothing with lowest variability across parcellations. These findings are further explained with reference to the number of resels and effective smoothing in section 3.5. Finally, while present across all brain areas, the variability between correlation maps is the highest within association cortices, primary motor, and insular cortex.

**Figure 3.**
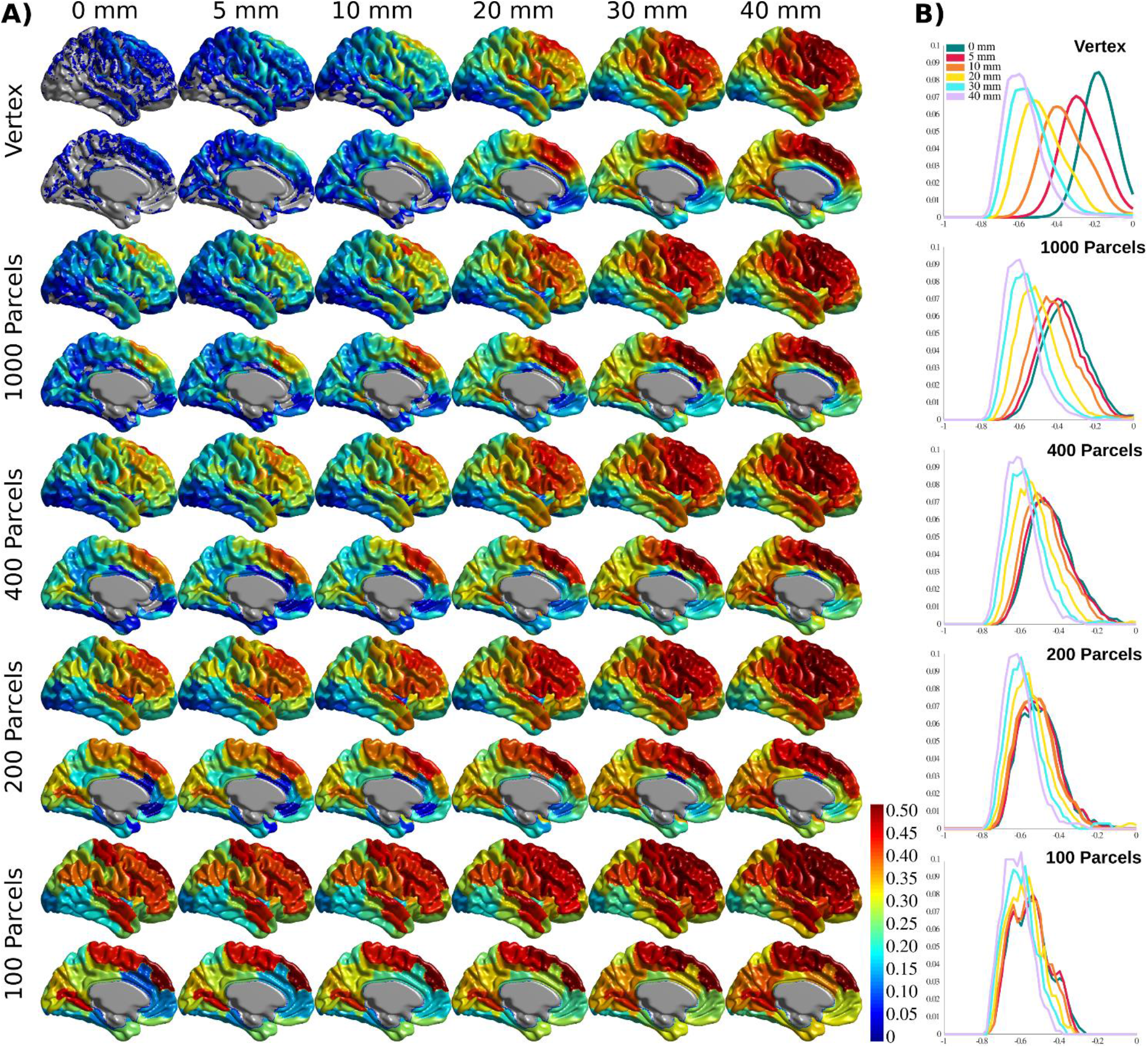
Cortical thickness variance explained by age (*r*^2^). **A)** Cortical thickness variance explained by age (*r*^2^) for each vertex/parcel across smoothing/parcellation conditions. **B)** Histograms for correlation values ® for each parcellation conditions, grouped by smoothing level.

### 3.4 brain age prediction based on cortical thickness

For age prediction, vertex-level data outperformed all parcellation-based data using the same (or a smaller) number of principal components as predictive features. The accuracy was also higher for lower smoothing kernel size. However, this effect was more pronounced for FWHMs greater than 10mm, and the results for FWHM values of 0, 5, and 10 mm showed a very similar performance in the vertex-level analysis. A similar pattern was present within each parcellation level. The best performing models (i.e. 0 and 5 mm smoothed vertex-wise), reach their minimum error using the first 20-30 principal components as features in the prediction model (i.e. a sample to feature ratio of 28-18). The pattern was similar for MAE and correlation between predicted age and chronological age (Supplementary Figure 2 and 3).

### 3.5 The relationship between prediction and association

As expected, there was a negative relationship between the overall correlation between age and cortical thickness across brain regions (measured by median *r*^2^) and the number of resels within each condition (Figure 5-A). Interestingly, we found a positive association between the number of resels and the overall ability of cortical thickness features to explain the variance of chronological age (as measured by adjusted *R*^2^ of the linear model) shown in Figure 5-B. These results suggest that the higher number of resels results in lower correlation values, but since resels are independent based on their relationship with age, they can explain different modes of chronological age within the population (hence the higher adjusted *R*^2^), whereas, in conditions with lower resel numbers (i.e. higher smoothing and larger parcels) the correlation values are higher but homogenous across the brain and therefore explain a lower proportion of the age variance.

Finally, there was a strong linear relationship between (i) the overall variance explained (adjusted *R*^2^) using a linear model with age as dependent variable and PCs as independent variable and (ii) the predictive performance of the linear regression model, with a bias due to overfitting in the linear model (Figure 5-C). Figure 5-D shows the overfitting bias of the adjusted *R*^2^ compared to the cross-validated prediction, as a function of the number of features in the model. Taken together, these results explain the opposing directions between correlation results and prediction accuracy across parcellation and smoothing conditions.

### 3.6 The effect of smoothing and parcellation on the estimation of brain age delta

In this section, we present **δ_2_** age prediction accuracy results with 10-fold cross validation. The prediction accuracy based on the modified **δ_2_** is presented in Figure 6. One of the main assumptions in age prediction studies is that delta age measured in different studies using different processing parameters are similar and can be interpreted as the same measure. We have examined the relationship between the optimal **δ_2_** across different parcellations and smoothing kernels (Figure 7). These results demonstrate the degree of sensitivity of **δ_2_** as a function of our choice for parcellation and smoothing kernel. While there is high correlation for large smoothing kernels (20-40 mm) as well as lower number of parcels, these conditions have the lowest prediction accuracies. The correlations between these conditions and higher accuracy conditions (i.e. vertex-wise and 1000 parcels with 0-10 mm smoothing) are lower (*r*~0.55). See the results for **δ_1_** in the Supplementary Figure 4.

**Figure 4.**
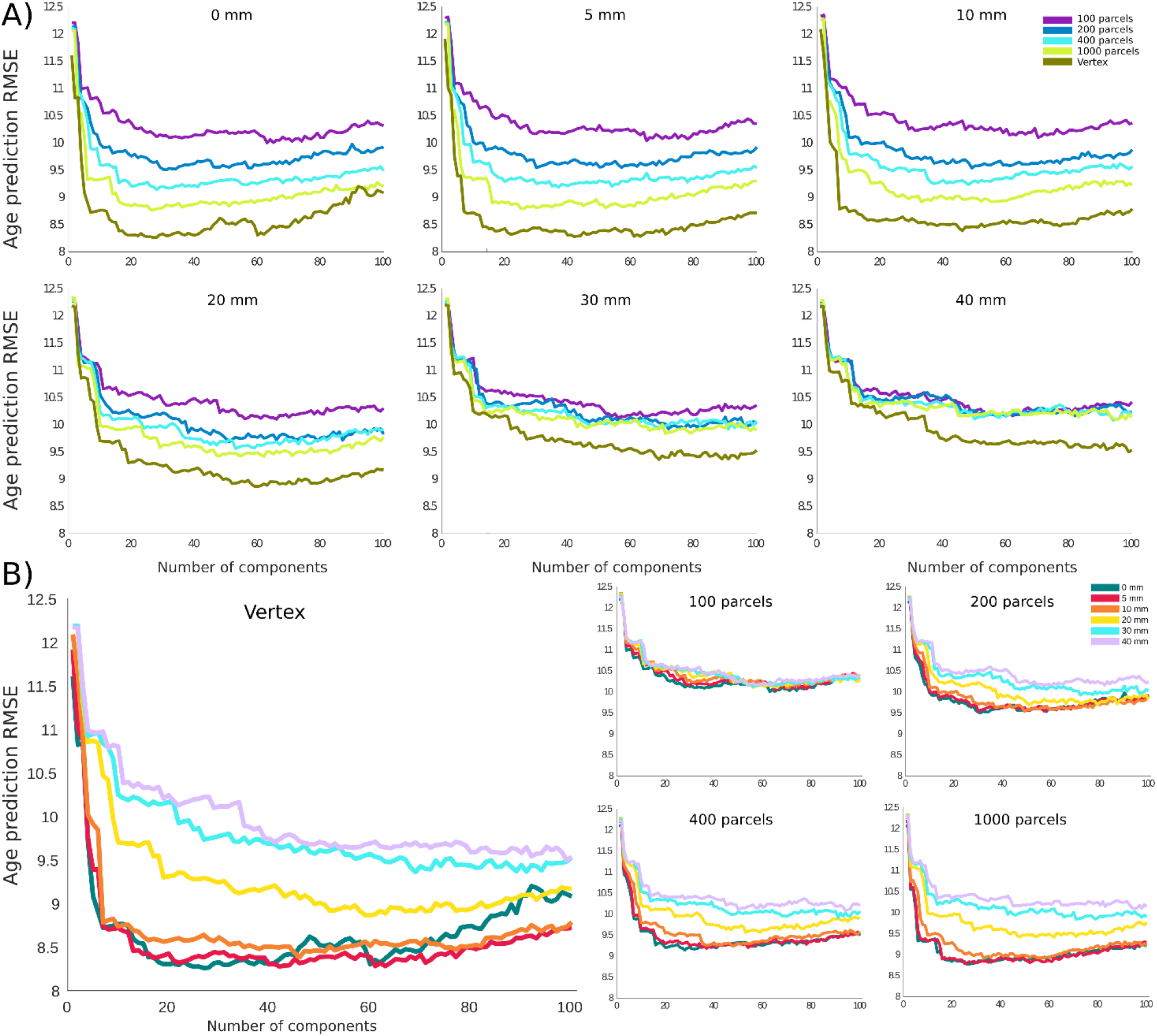
Root mean square error (RMSE) for age prediction as a function of number of principal components included as features in the predictive model. **A)** Results grouped together based on the smoothing level. **B)** Results grouped together based on the parcellation resolution.

## 4 Discussion

In this article, we compared the effect of different smoothing and parcellation on associations between cortical thickness and chronological age as well as brain age prediction accuracy. We showed that the optimal choice for association analysis might indeed undermine age prediction accuracy, and vice versa. We further investigated this relationship and demonstrated the underlying differences that lead to this trade-off between the two analyses. Finally, we examined the effect of smoothing and parcellation on delta age estimation and showed that the initial smoothing and parcellation choices can change the delta estimation which in turn will affect any downstream analysis.

We used brain association with age and brain age prediction as our target analyses, since age is used as the main variable of interest or at least a confounding variable in most neuroimaging studies. We used cortical thickness as the main measure of interest. Due to the wide availability of T1-weighted MRI in research and clinical settings, cortical thickness is a suitable measure which has been widely used to study brain anatomy in general (Toga, 2015), and more specifically, brain aging and predicting brain age (Groves et al., 2012; Kandel et al., 2013; Liem et al., 2017; Wang & Pham, 2011). Finally, our results are presented based on a sample size of N~600 which is a common sample size for publicly available datasets in the field of neuroimaging.

Given the limited number of subjects in neuroimaging studies compared to potential features (number of vertices/voxels), most prediction studies apply dimension reduction as an initial step. We used PCA for dimension reduction of the cortical thickness data. Due to its simplicity and interpretability, PCA has been widely used in the brain age prediction literature. Furthermore, we employed linear regression with cross-validation as our prediction model (Smith et al., 2019a). As expected, we observed an initial drop in the prediction error, followed by a plateau/increase in the error as the sample to feature ratio increases (Hastie et al., 2009). At each parcellation level, the accuracy drops with increased smoothing, and for each smoothing level, the accuracy decreases with larger parcels/regions.

It is commonplace for neuroimaging studies to use smoothing and parcellation as the first step of their analysis to achieve higher statistical power with reducing the individual variability within the data. Furthermore, with increased availability of public neuroimaging datasets, it is commonplace to release a preprocessed version of the data with a fixed smoothing level and averaged based on a given parcellation. Many research groups in the field use preprocessed and parcellation-based data releases as the starting point for their analyses. In fact, in many cases, the raw data is not publicly distributed, and the preprocessed parcellated data is the only version of data available. For example, some of the most influential public datasets in the field of neuroimaging such as Adolescent Brain Cognitive Development (ABCD, for details see https://nda.nih.gov/abcd) Study and UKBiobank (for details see https://www.ukbiobank.ac.uk) provide cortical thickness data using Desikan-Killiany-Tourville parcellations (Klein & Tourville, 2012) with 62 regions (smoothing varies across studies) as one of their pre-calculated measures. Our findings can help provide a guide to interpret these available measures and shed light on the effect of these preselected parameters/parcellation when applied in aging studies.

Higher correlation values across brain regions (as a result of smoothing) can be explained by increased signal to noise ratio and reduced individual variability (Figure 2). The effect of smoothing on brain related associations has previously been studied (Lerch & Evans, 2005). Indeed, Zhao and colleagues propose smoothing as a scaling dimension which needs optimization for any given target analysis (Zhao et al., 2013). The effect of parcellation on brain association has been addressed in several studies. However, the optimal parcellation level is still an open question dependent on the specific case of interest (Eickhoff et al., 2018). Here, we showed that parcellation level has a similar impact, by reducing variability, using both CV (Figure 2) and number of resels (Figure 1).

In neuroimaging, smoothing and parcellations are generally studied separately. In this study, we used a unified metric to directly compare the effect of smoothing and parcellation. Using resel numbers and variance explained in the model, we have calculated common measures for both association and prediction results. Our results show that with increased smoothing and larger parcels (i.e. lower number of resels), cortical thickness variability reduces. This will remove inter-individual differences across brain regions and result in higher associations between cortical thickness and aging (Figure 5-A). However, while this improves the regional correlation with age, most of this general trend can be captured in a few PCs (mainly the first component) and the rest of the PCs do not explain the remaining variance of age. On the other hand, this relationship is reversed in the conditions with higher resel numbers (i.e. lower smoothing and higher spatial resolutions). While in these cases higher regional variability results in lower correlation with age, the age related associations capture different portions of age variance in different PCs and overall they have a higher adjusted *R*^2^ (Figure 5-B). There was a consistent bias in the adjusted *R*^2^ across conditions (Figure 5-C and 5-D), however, the effects remained similar after removing the overfitting with cross-validation. Altogether, these analyses explain the seeming opposite direction of correlation values and prediction accuracies for different smoothing/parcellation levels in section 3.3 and 3.4.

**Figure 5.**
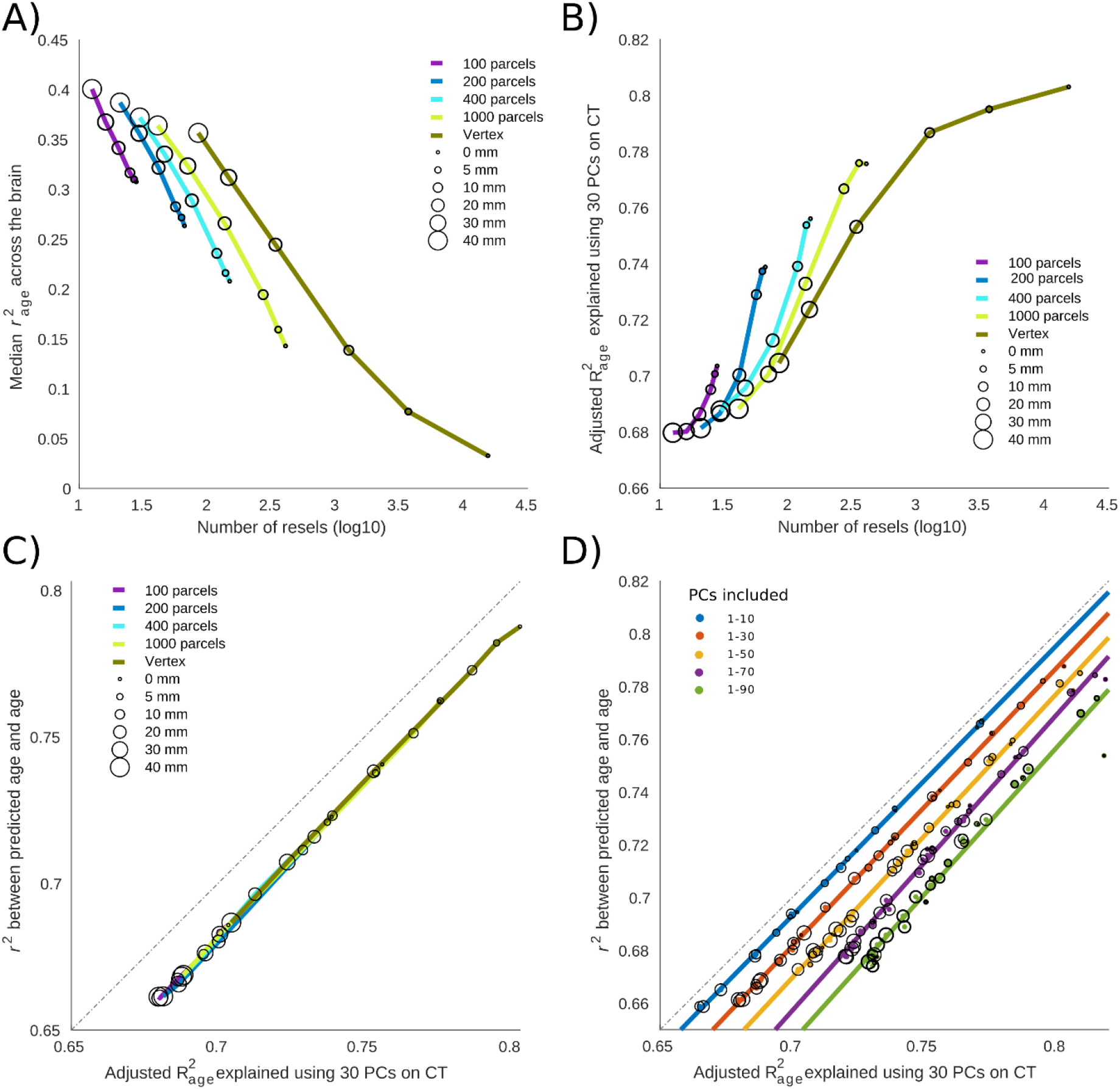
The relationship between cortical thickness association with age versus brain age prediction. **A)** Median variance explained of cortical thickness across the brain. The results are grouped based on the parcellation. Circles represent the something level within each parcellation. **B)** Total variance explained of age by the first 30 principal components (PCs) of cortical thickness as independent variables. **C)** The relationship between age prediction accuracy and total variance explained of age. In the case of prediction, the first PCs are used as predictive features alongside cross validation to prevent overfitting. The total variance explained of age is the same as depicted in B. **D)** The overfitting bias of linear model compared to the same model used with cross validation. As expected, a higher number of predictive features results in higher level of overfitting bias.

**Figure 6.**
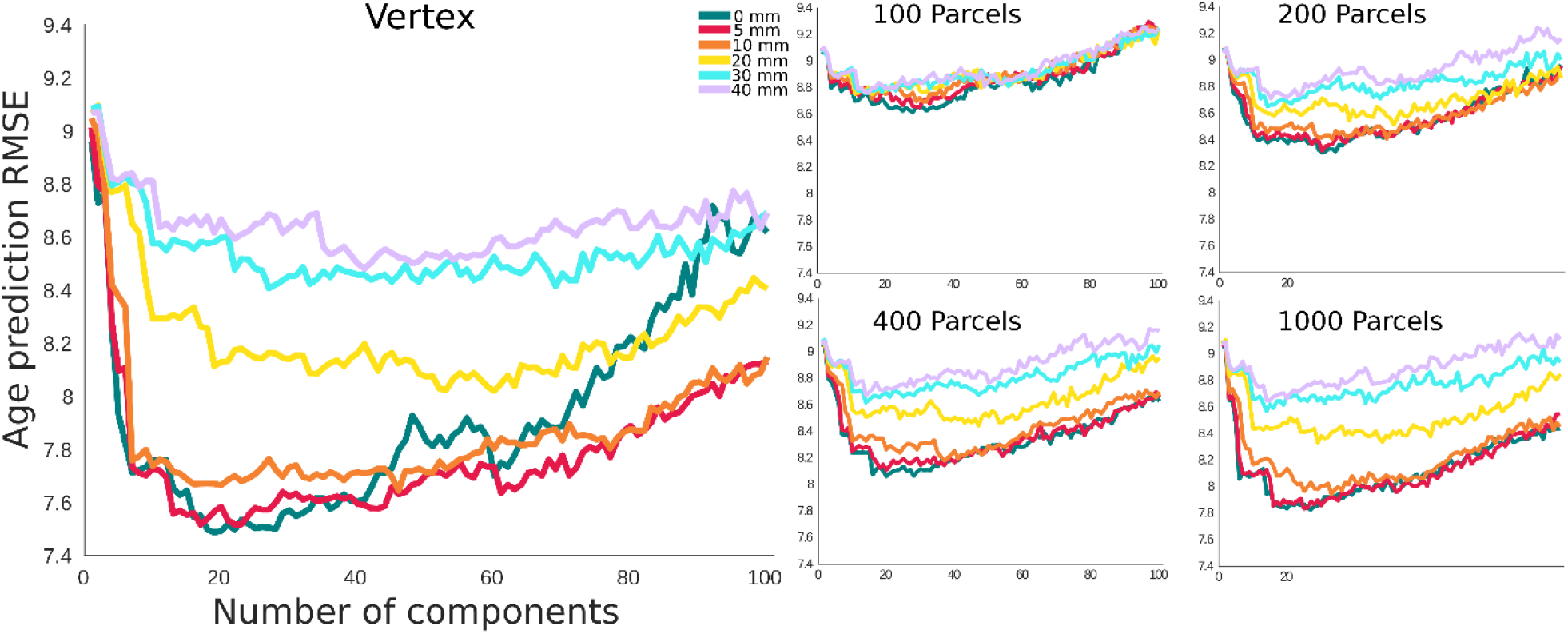
Root mean square error (RMSE) for Age prediction with **δ_2_** as the error term. The x axis shows the number of principal components included as features. The results are grouped based on the parcellation resolution.

**Figure 7.**
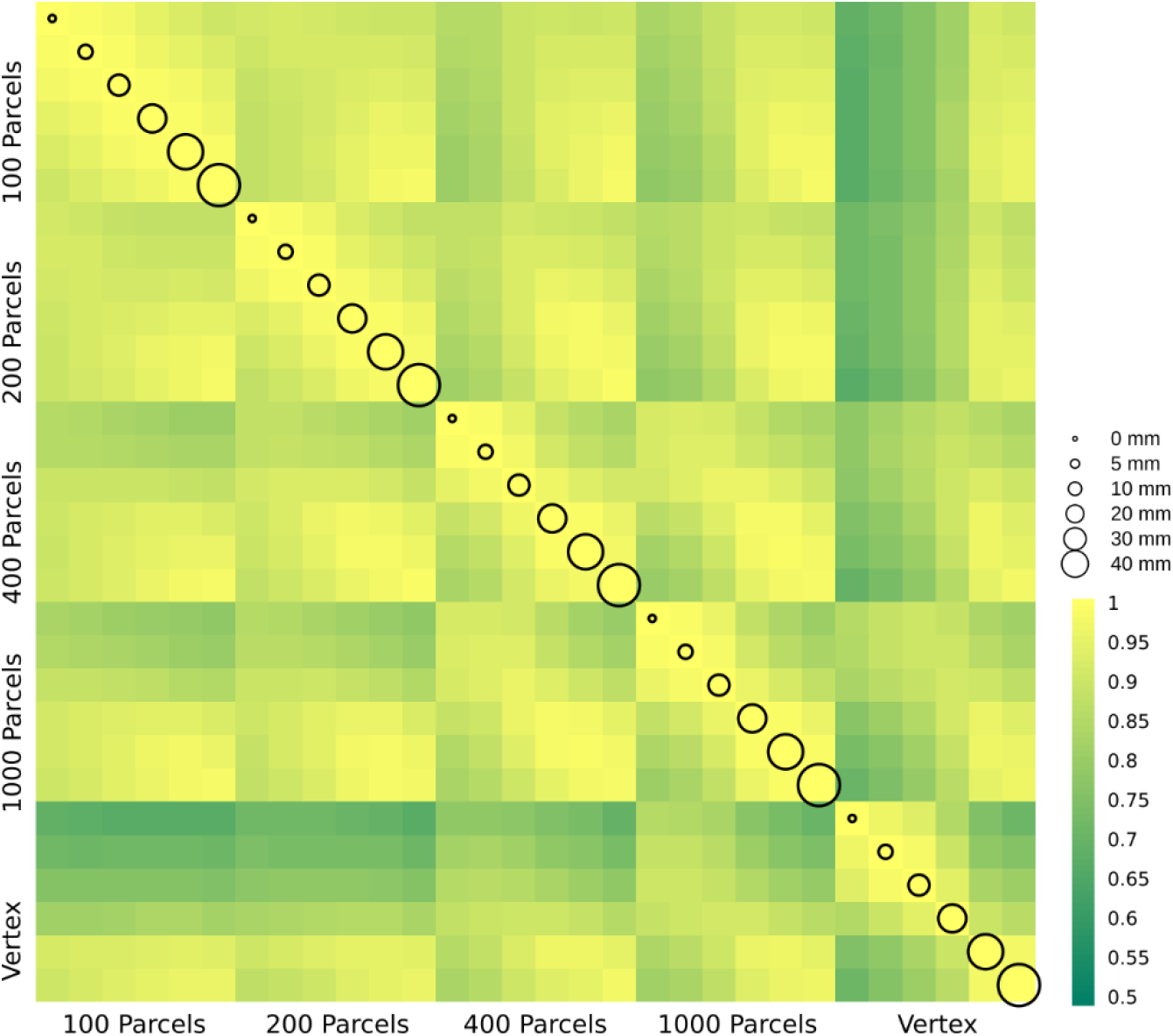
**δ_2_** age prediction error. The correlation between delta age (as measured by **δ_2_**) across parcellation resolutions (x and y axis labels) and smoothing kernels (represented by circle size).

Brain age studies investigate the relationship between and other phenotypes, using a given smoothing and vertex/parcellation resolution as their initial step (James H. Cole & Franke, 2017). However, the effect of the preprocessing condition on estimation is not studied. In the current manuscript, we found a range of associations (0.5-1) between **δ_2_**s obtained in different conditions. These results suggest not only that each study needs to optimize their choice of the smoothing and parcellation level, but also when interpreting results from different studies in the field, these parameters should be considered.

One of the main limitations of the current study is the number of subjects (N~600), particularly given that their age spans across 70 years. This leads to overfitting as the number of features increase. In fact, for vertex-wise prediction (with 0 mm smoothing), the first 30 PCs only explain 20 % of the variability in the data. This number is around 40% for 10 mm smoothing. In comparison, the first 30 PCs for 100 parcels explain 80 and 90% of the variance of the cortical thickness data for 0 mm and 40 mm smoothing levels, respectively (Supplementary Figure 5). Given the higher performance of the vertex-wise PCs at 0-10 mm smoothing, it is likely that with a larger sample size and increased sample to feature ratio, the accuracy can be further improved. It should be noted that in each case the variance explained corresponds to the total variability for the corresponding smoothing and parcellation condition. Another limitation in the current study is the use of functionally driven Schaefer parcellations. While this does not automatically suggest a disadvantage, multi-resolution anatomically driven parcellations have the theoretical advantage of a more relevant initial feature space for cortical thickness studies. Finally, CamCAN data used in our study is cross-sectional. This potentially decreases the detection power of our study, since we can only estimate the effect of time between subjects with individual variability as part of the measurement, whereas a longitudinal dataset can decrease variability by estimating the effect of aging within subjects.

Traditionally, neuroimaging studies have targeted brain related associations with a given phenotype/symptom or the statistical differences between different groups for a given brain region, followed up with the association of these differences with a given biological or behavioral variable of interest. More recently, there has been an ongoing conversation in the field towards prediction as an alternative approach. Along the same line, the field of brain aging, has pursued age related associations as well as age prediction. The relationship between the two approaches is often taken for granted (since in ideal settings, i.e. large sample size and low inter-individual variability or noise levels, the results would be equivalent) and ignored in practice. In this study, we have directly addressed both age association and prediction as a function of smoothing and parcellation levels. Within our sample size, we found an inverse relationship between regional age related associations and brain age prediction accuracy as a function of smoothing and parcellation level, highlighting the importance of the parameter selection based on the goal of the study.

## 6 Author Contributions

YZ contributed to the study plan, analysed the data, and wrote the manuscript. AE contributed to the study plan and revision of the manuscript.

## 8 Acknowledgments

Data collection and sharing for this project was provided by the Cambridge Centre for Ageing and Neuroscience (CamCAN). CamCAN funding was provided by the UK Biotechnology and Biological Sciences Research Council (grant number BB/H008217/1), together with support from the UK Medical Research Council and University of Cambridge, UK. Authors thank Compute Canada (https://www.computecanada.ca/home) for the usage of the computing resources in the current work.

Authors thank Dr. Lindsay B Lewis for providing the surface parcellations registered to the MNI surface space as well as Drs Mahsa Dadar and Filip Morys for their inputs and comments regarding data analysis and the final manuscript.

## 9 Data Availability Statement

CamCAN dataset can be accessed after submitting an online application at https://camcan-archive.mrc-cbu.cam.ac.uk/dataaccess/datarequest.php and accepting the use agreement. Detailed information available at https://www.cam-can.org.

## Reference

Booth, T., Starr, J. M., & Deary, I. (2013). Modeling multisystem biological risk in later life: Allostatic load in the lothian birth cohort study 1936. American Journal of Human Biology, 25(4), 538–543. https://doi.org/10.1002/ajhb.22406

Bzdok, D., Varoquaux, G., & Steyerberg, E. W. (2020). Prediction, not association, paves the road to precision medicine. In JAMA Psychiatry. https://doi.org/10.1001/jamapsychiatry.2020.2549

Chung, M., Worsley, K., Paus, T., Robbins, S., Evans, A. C., Taylor, J., Giedd, J. N., & Rapoport, J. L. (2002). Tensor-based surface morphometry. In University of Wisconsin (Issue 1049). http://www.stat.wisc.edu/~mchung

Cole, J. H., Ritchie, S. J., Bastin, M. E., Valdés Hernández, M. C., Muñoz Maniega, S., Royle, N., Corley, J., Pattie, A., Harris, S. E., Zhang, Q., Wray, N. R., Redmond, P., Marioni, R. E., Starr, J. M., Cox, S. R., Wardlaw, J. M., Sharp, D. J., & Deary, I. J. (2018). Brain age predicts mortality. Molecular Psychiatry, 23(5), 1385–1392. https://doi.org/10.1038/mp.2017.62

Cole, James H., & Franke, K. (2017). Predicting Age Using Neuroimaging: Innovative Brain Ageing Biomarkers. In Trends in Neurosciences (Vol. 40, Issue 12, pp. 681–690). Elsevier Ltd. https://doi.org/10.1016/j.tins.2017.10.001

Collins, D. L., Neelin, P., Peters, T. M., & Evans, A. C. (1994). Automatic 3d intersubject registration of mr volumetric data in standardized talairach space. Journal of Computer Assisted Tomography, 18(2), 192–205. https://doi.org/10.1097/00004728-199403000-00005

Curiati, P. K., Tamashiro, J. H., Squarzoni, P., Duran, F. L. S., Santos, L. C., Wajngarten, M., Leite, C., Vallada, H., Menezes, P. R., Scazufca, M., Busatto, G. F., & Alves, T. C. T. F. (2009). Brain structural variability due to aging and gender in cognitively healthy elders: Results from the São Paulo ageing and health study. American Journal of Neuroradiology, 30(10), 1850–1856. https://doi.org/10.3174/ajnr.A1727

Eickhoff, S. B., Yeo, B. T. T., & Genon, S. (2018). Imaging-based parcellations of the human brain. In Nature Reviews Neuroscience (Vol. 19, Issue 11, pp. 672–686). https://doi.org/10.1038/s41583-018-0071-7

Franke, K., & Gaser, C. (2019a). Ten years of brainage as a neuroimaging biomarker of brain aging: What insights have we gained? In Frontiers in Neurology (Vol. 10, Issue JUL). Frontiers Media S.A. https://doi.org/10.3389/fneur.2019.00789

Franke, K., & Gaser, C. (2019b). Ten years of brainage as a neuroimaging biomarker of brain aging: What insights have we gained? In Frontiers in Neurology (Vol. 10, Issue JUL). https://doi.org/10.3389/fneur.2019.00789

Franke, K., Luders, E., May, A., Wilke, M., & Gaser, C. (2012). Brain maturation: Predicting individual BrainAGE in children and adolescents using structural MRI. NeuroImage, 63(3), 1305–1312. https://doi.org/10.1016/j.neuroimage.2012.08.001

Gaser, C., Franke, K., Klöppel, S., Koutsouleris, N., & Sauer, H. (2013). BrainAGE in Mild Cognitive Impaired Patients: Predicting the Conversion to Alzheimer’s Disease. PLoS ONE, 8(6). https://doi.org/10.1371/journal.pone.0067346

Gorgolewski, K., Tambini, A., Durnez, J., Sochat, V., Wexler, J., & Poldrack, R. (2016). Evaluation of full brain parcellation schemes using the NeuroVault database of statistical maps. Organisation for Human Brain Mapping 2016 Annual Meeting, 2201. https://54.246.141.91/posters/6-1986

Groves, A. R., Smith, S. M., Fjell, A. M., Tamnes, C. K., Walhovd, K. B., Douaud, G., Woolrich, M. W., & Westlye, L. T. (2012). Benefits of multi-modal fusion analysis on a large-scale dataset: Life-span patterns of inter-subject variability in cortical morphometry and white matter microstructure. NeuroImage, 63(1), 365–380. https://doi.org/10.1016/j.neuroimage.2012.06.038

Hastie, T., Tibshirani, R., & Friedman, J. (2009). Springer Series in Statistics The Elements of Statistical Learning Data Mining, Inference, and Prediction. https://books.google.ca/books?hl=en&lr=&id=tVIjmNS3Ob8C&oi=fnd&pg=PR13&dq=trevor+hastie++book&ots=EOBcP9J5X5&sig=w9Dod2i1zZD9tkzSKn0TPwDs1UE

Hayasaka, S., Phan, K. L., Liberzon, I., Worsley, K. J., & Nichols, T. E. (2004). Nonstationary cluster-size inference with random field and permutation methods. NeuroImage, 22(2), 676–687. https://doi.org/10.1016/j.neuroimage.2004.01.041

Horvath, S. (2013). DNA methylation age of human tissues and cell types. Genome Biology, 14(10). https://doi.org/10.1186/gb-2013-14-10-r115

Hu, S., Chao, H. H. A., Zhang, S., Ide, J. S., & Li, C. S. R. (2014). Changes in cerebral morphometry and amplitude of low-frequency fluctuations of BOLD signals during healthy aging: Correlation with inhibitory control. Brain Structure and Function, 219(3), 983–994. https://doi.org/10.1007/s00429-013-0548-0

Jones, S. E., Buchbinder, B. R., & Aharon, I. (2000). Three-dimensional mapping of cortical thickness using Laplace’s equation. Human Brain Mapping, 11(1), 12–32. https://doi.org/10.1002/1097-0193(200009)11:1<12::AID-HBM20>3.0.CO;2-K

Kabani, N., Le Goualher, G., Macdonald, D., & Evans, A. C. (2001). Measurement of Cortical Thickness Using an Automated 3-D Algorithm: A Validation Study. Elsevier. https://doi.org/10.1006/nimg.2000.0652

Kandel, B. M., Wolk, D. A., Gee, J. C., & Avants, B. (2013). Predicting cognitive data from medical images using sparse linear regression. Lecture Notes in Computer Science (Including Subseries Lecture Notes in Artificial Intelligence and Lecture Notes in Bioinformatics), 7917 LNCS, 86–97. https://doi.org/10.1007/978-3-642-38868-2_8

Kaufmann, T., van der Meer, D., Doan, N. T., Schwarz, E., Lund, M. J., Agartz, I., Alnæs, D., Barch, M., Baur-Streubel, R., Bertolino, A., Bettella, F., Beyer, M. K., Bøen, E., Borgwardt, S., Brandt, C. L., Buitelaar, J., Celius, E. G., Cervenka, S., Conzelmann, A., … Westlye, L. T. (2019). Common brain disorders are associated with heritable patterns of apparent aging of the brain. Nature Neuroscience, 22(10), 1617–1623. https://doi.org/10.1038/s41593-019-0471-7

Khundrakpam, B. S., Tohka, J., Evans, A. C., Ball, W. S., Byars, A. W., Schapiro, M., Bommer, W., Carr, A., German, A., Dunn, S., Rivkin, M. J., Waber, D., Mulkern, R., Vajapeyam, S., Chiverton, A., Davis, P., Koo, J., Marmor, J., Mrakotsky, C., … O’Neill, J. (2015). Prediction of brain maturity based on cortical thickness at different spatial resolutions. NeuroImage, 111, 350–359. https://doi.org/10.1016/j.neuroimage.2015.02.046

Kim, J., Singh, V., Jun, K. L., Lerch, J., Ad-Dab’bagh, Y., MacDonald, D., Jong, M. L., Kim, S. I., & Evans, A. C. (2005). Automated 3-D extraction and evaluation of the inner and outer cortical surfaces using a Laplacian map and partial volume effect classification. NeuroImage, 27(1), 210–221. https://doi.org/10.1016/j.neuroimage.2005.03.036

Klein, A., & Tourville, J. (2012). 101 labeled brain images and a consistent human cortical labeling protocol. Frontiers in Neuroscience, DEC. https://doi.org/10.3389/fnins.2012.00171

Le, T. T., Kuplicki, R. T., McKinney, B. A., Yeh, H.-W., Thompson, W. K., & Paulus, M. P. (2018). A Nonlinear Simulation Framework Supports Adjusting for Age When Analyzing BrainAGE. Frontiers in Aging Neuroscience, 10. https://doi.org/10.3389/fnagi.2018.00317

Lemaître, H., Crivello, F., Grassiot, B., Alpérovitch, A., Tzourio, C., & Mazoyer, B. (2005). Age- and sex-related effects on the neuroanatomy of healthy elderly. NeuroImage, 26(3), 900–911. https://doi.org/10.1016/j.neuroimage.2005.02.042

Lerch, J. P., & Evans, A. C. (2005). Cortical thickness analysis examined through power analysis and a population simulation. NeuroImage, 24(1), 163–173. https://doi.org/10.1016/j.neuroimage.2004.07.045

Lerch, J. P., Worsley, K., Shaw, W. P., Greenstein, D. K., Lenroot, R. K., Giedd, J., & Evans, A. C. (2006). Mapping anatomical correlations across cerebral cortex (MACACC) using cortical thickness from MRI. NeuroImage, 31(3), 993–1003. https://doi.org/10.1016/j.neuroimage.2006.01.042

Lewis, J. D., Fonov, V. S., Collins, D. L., Evans, A. C., & Tohka, J. (2019). Cortical and subcortical T1 white/gray contrast, chronological age, and cognitive performance. NeuroImage, 196, 276–288. https://doi.org/10.1016/j.neuroimage.2019.04.022

Lewis, L. B., Lepage, C. Y., & Evans, A. C. (2019). An extended MSM surface registration pipeline to bridge atlases across the MNI and the FS/HCP worlds. Annual Meeting of the Organization for Human Brain Mapping. https://ww5.aievolution.com/hbm1901/index.cfm?do=abs.viewAbs&abs=1243

Liang, H., Zhang, F., & Niu, X. (2019). Investigating systematic bias in brain age estimation with application to post-traumatic stress disorders. Human Brain Mapping, 40(11), 3143–3152. https://doi.org/10.1002/hbm.24588

Liem, F., Varoquaux, G., Kynast, J., Beyer, F., Kharabian Masouleh, S., Huntenburg, J. M., Lampe, L., Rahim, M., Abraham, A., Craddock, R. C., Riedel-Heller, S., Luck, T., Loeffler, M., Schroeter, M. L., Witte, A. V., Villringer, A., & Margulies, D. S. (2017). Predicting brain-age from multimodal imaging data captures cognitive impairment. NeuroImage, 148, 179–188. https://doi.org/10.1016/j.neuroimage.2016.11.005

López-Otín, C., Blasco, M. A., Partridge, L., Serrano, M., & Kroemer, G. (2013). The hallmarks of aging. In Cell (Vol. 153, Issue 6, p. 1194). Cell Press. https://doi.org/10.1016/j.cell.2013.05.039

MacDonald, D., Kabani, N., Avis, D., & Evans, A. C. (2000). Automated 3-D extraction of inner and outer surfaces of cerebral cortex from MRI. NeuroImage, 12(3), 340–356. https://doi.org/10.1006/nimg.1999.0534

Salehi, M., Greene, A. S., Karbasi, A., Shen, X., Scheinost, D., & Constable, R. T. (2020). There is no single functional atlas even for a single individual: Functional parcel definitions change with task. NeuroImage, 208. https://doi.org/10.1016/j.neuroimage.2019.116366

Schaefer, A., Kong, R., Gordon, E. M., Laumann, T. O., Zuo, X.-N., Holmes, A. J., Eickhoff, S. B., & Yeo, B. T. T. (2018). Local-Global Parcellation of the Human Cerebral Cortex from Intrinsic Functional Connectivity MRI. Cerebral Cortex, 28(9), 3095–3114. https://doi.org/10.1093/cercor/bhx179

Shafto, M. A., Tyler, L. K., Dixon, M., Taylor, J. R., Rowe, J. B., Cusack, R., Calder, A. J., Marslen-Wilson, W. D., Duncan, J., Dalgleish, T., Henson, R. N., Brayne, C., Bullmore, E., Campbell, K., Cheung, T., Davis, S., Geerligs, L., Kievit, R., McCarrey, A., … Matthews, F. E. (2014). The Cambridge Centre for Ageing and Neuroscience (Cam-CAN) study protocol: A cross-sectional, lifespan, multidisciplinary examination of healthy cognitive ageing. BMC Neurology, 14(1). https://doi.org/10.1186/s12883-014-0204-1

Sled, J. G., Zijdenbos, A. P., & Evans, A. C. (1998). A nonparametric method for automatic correction of intensity nonuniformity in mri data. IEEE Transactions on Medical Imaging, 17(1), 87–97. https://doi.org/10.1109/42.668698

Smith, S. M., Vidaurre, D., Alfaro-Almagro, F., Nichols, T. E., & Miller, K. L. (2019a). Estimation of Brain Age Delta from Brain Imaging. In bioRxiv. https://doi.org/10.1101/560151

Smith, S. M., Vidaurre, D., Alfaro-Almagro, F., Nichols, T. E., & Miller, K. L. (2019b). Estimation of brain age delta from brain imaging. NeuroImage, 200, 528–539. https://doi.org/10.1016/j.neuroimage.2019.06.017

Storsve, A. B., Fjell, A. M., Tamnes, C. K., Westlye, L. T., Overbye, K., Aasland, H. W., & Walhovd, K. B. (2014). Differential longitudinal changes in cortical thickness, surface area and volume across the adult life span: Regions of accelerating and decelerating change. Journal of Neuroscience, 34(25), 8488–8498. https://doi.org/10.1523/JNEUROSCI.0391-14.2014

Takahashi, R., Ishii, K., Kakigi, T., & Yokoyama, K. (2011). Gender and age differences in normal adult human brain: Voxel-based morphometric study. Human Brain Mapping, 32(7), 1050–1058. https://doi.org/10.1002/hbm.21088

Taylor, J. R., Williams, N., Cusack, R., Auer, T., Shafto, M. A., Dixon, M., Tyler, L. K., Cam-CAN, & Henson, R. N. (2017). The Cambridge Centre for Ageing and Neuroscience (Cam-CAN) data repository: Structural and functional MRI, MEG, and cognitive data from a cross-sectional adult lifespan sample. NeuroImage, 144, 262–269. https://doi.org/10.1016/j.neuroimage.2015.09.018

Thomas Yeo, B. T., Krienen, F. M., Sepulcre, J., Sabuncu, M. R., Lashkari, D., Hollinshead, M., Roffman, J. L., Smoller, J. W., Zöllei, L., Polimeni, J. R., Fisch, B., Liu, H., & Buckner, R. L. (2011). The organization of the human cerebral cortex estimated by intrinsic functional connectivity. Journal of Neurophysiology, 106(3), 1125–1165. https://doi.org/10.1152/jn.00338.2011

Toga, A. W. (2015). Brain Mapping: An Encyclopedic Reference. In Brain Mapping: An Encyclopedic Reference (Vols. 1-3). https://doi.org/10.1016/C2011-1-07037-6

Tohka, J., Zijdenbos, A., & Evans, A. (2004). Fast and robust parameter estimation for statistical partial volume models in brain MRI. NeuroImage, 23(1), 84–97. https://doi.org/10.1016/j.neuroimage.2004.05.007

von Economo, CF; Koskinas, G. (1927). Die Cytoarchitektonik der Hirnrinde des erwachsenen Menschen. Journal of Anatomy, 61, 264–266. https://scholar.google.com/scholar?q=Von+Economo+C%2C+Koskinas+G.+Die+Cytoarchitektonik+der+Hirnrinde+des+erwachsenen+Menschen.+Berlin%3A+Springer%3B+1925.

Wang, B., & Pham, T. D. (2011). MRI-based age prediction using hidden Markov models. Journal of Neuroscience Methods, 199(1), 140–145. https://doi.org/10.1016/j.jneumeth.2011.04.022

Worsley, K. J. (1996). An unbiased estimator for the roughness of a multivariate Gaussian random eld 1 Model. Pdfs.Semanticscholar.Org, 1–5. https://pdfs.semanticscholar.org/4159/a64da50863a945c8af38c42f8b09487a985b.pdf

Worsley, K. J., Andermann, M., Koulis, T., MacDonald, D., & Evans, A. C. (1999). Detecting changes in nonisotropic images. Human Brain Mapping, 8(2–3), 98–101. https://doi.org/10.1002/(SICI)1097-0193(1999)8:2/3<98::AID-HBM5>3.0.CO;2-F

Worsley, K. J., Evans, A. C., Marrett, S., & Neelin, P. (1992). A three-dimensional statistical analysis for CBF activation studies in human brain. Journal of Cerebral Blood Flow and Metabolism, 12(6), 900–918. https://doi.org/10.1038/jcbfm.1992.127

Yarkoni, T., & Westfall, J. (2017). Choosing Prediction Over Explanation in Psychology: Lessons From Machine Learning. Perspectives on Psychological Science, 12(6), 1100–1122. https://doi.org/10.1177/1745691617693393

Zhao, L., Boucher, M., Rosa-Neto, P., & Evans, A. C. (2013). Impact of scale space search on age- and gender-related changes in MRI-based cortical morphometry. Human Brain Mapping, 34(9), 2113–2128. https://doi.org/10.1002/hbm.22050

Ziegler, G., Dahnke, R., Jäncke, L., Yotter, R. A., May, A., & Gaser, C. (2012). Brain structural trajectories over the adult lifespan. Human Brain Mapping, 33(10), 2377–2389. https://doi.org/10.1002/hbm.21374

Zijdenbos, A. P., Forghani, R., & Evans, A. C. (2002). Automatic “pipeline” analysis of 3-D MRI data for clinical trials: Application to multiple sclerosis. IEEE Transactions on Medical Imaging, 21(10), 1280–1291. https://doi.org/10.1109/TMI.2002.806283

