## Supplementary Figures for "Association versus Prediction: the impact of cortical surface smoothing and parcellation on brain age"

### *Supplementary Material*

#### 1 Supplementary Figures

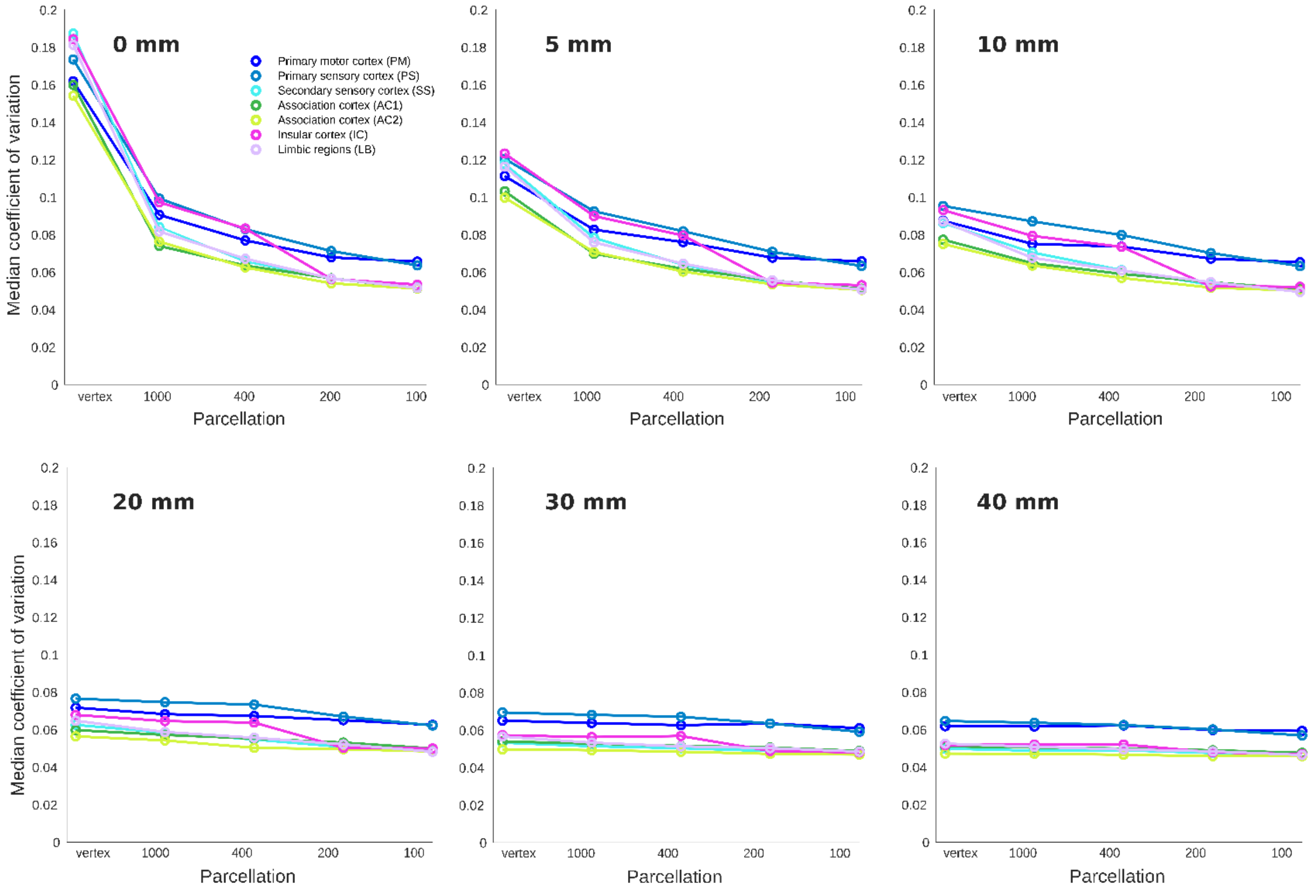


**Supplementary Figure 1.** Coefficient of variation (CV) shown for each cytoarchitectural region across parcellation and smoothing conditions.

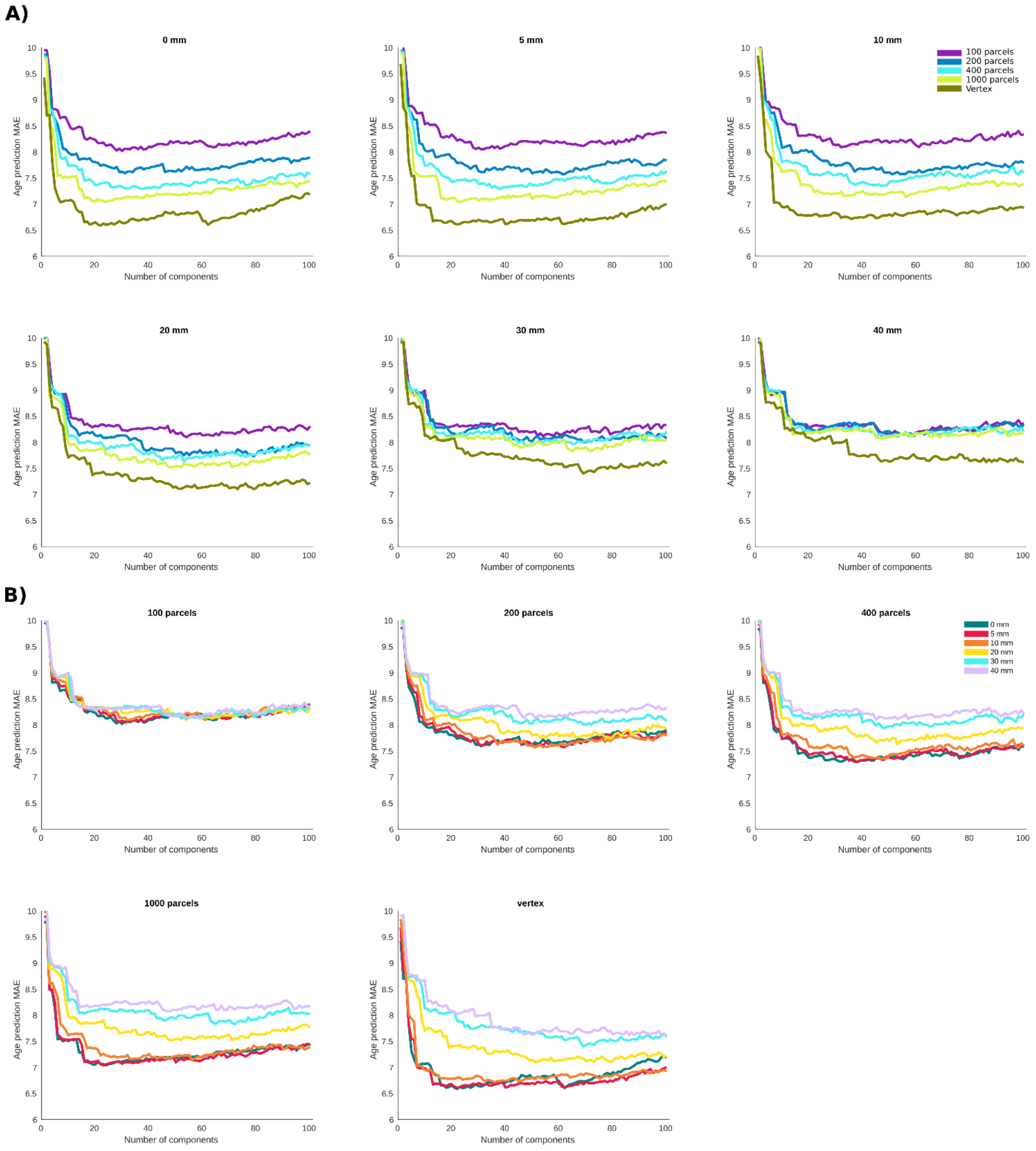


**Supplementary Figure 2.** Mean absolute error (MAE) for Age prediction as a function of number of principal components included as features in the predictive model. **A)** results grouped together based on the smoothing level. **B)** results grouped together based on the parcellation resolution.

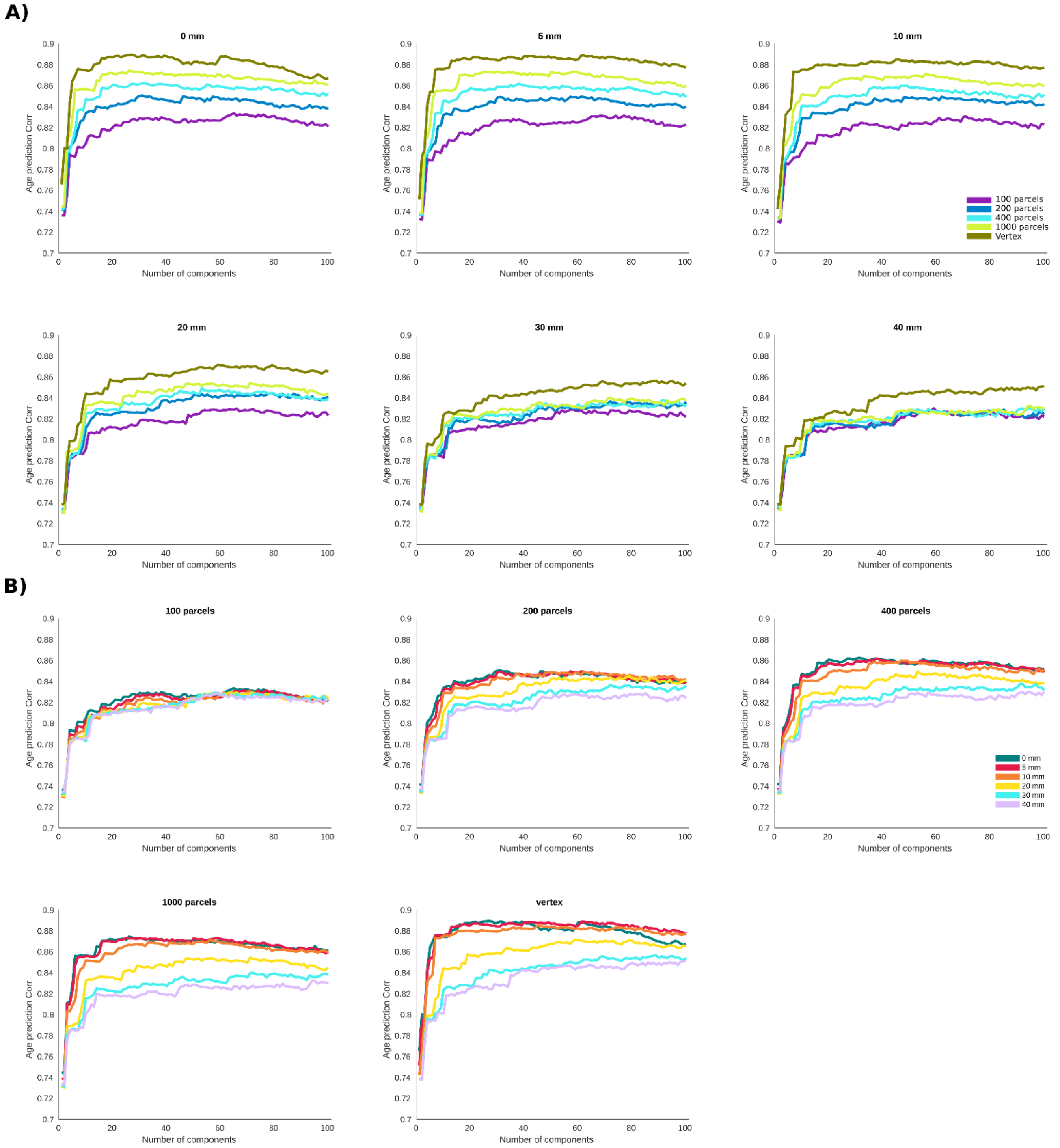


**Supplementary Figure 3.** Correlation between the predicted age and chronological age as a function of number of principal components included as features in the predictive model. **A)** results grouped together based on the smoothing level. **B)** results grouped together based on the parcellation resolution.

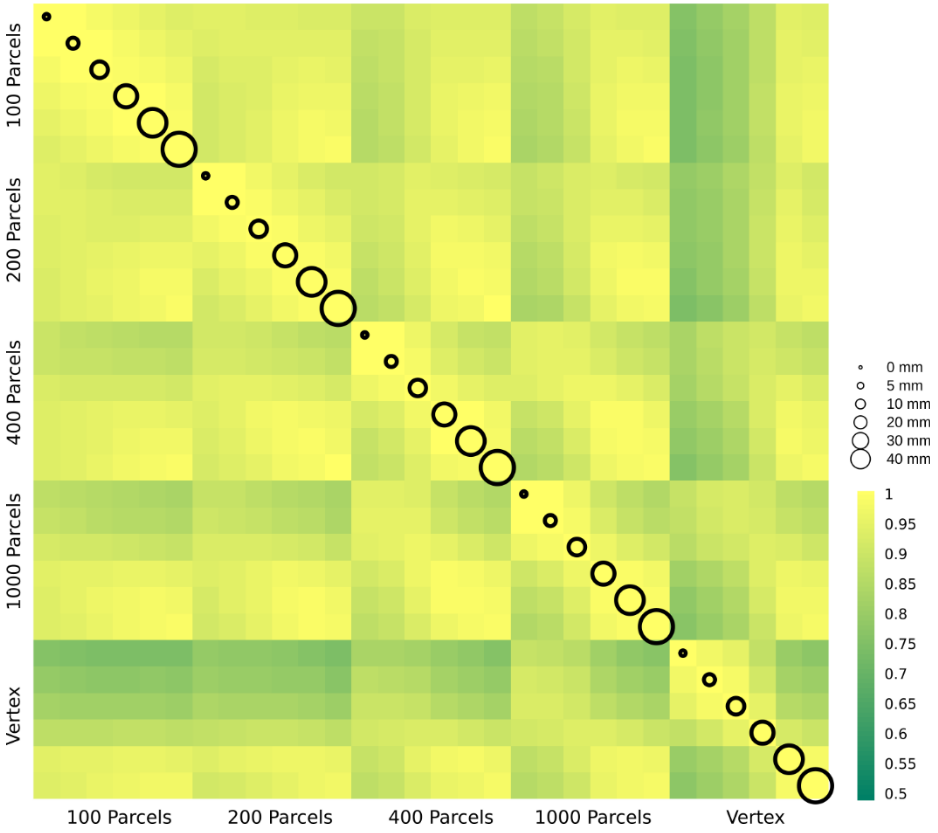


**Supplementary Figure 4.** $\delta_1$ age prediction error. The correlation between delta age (as measured by $\delta_1$) across parcellation resolutions (x axis labels) and smoothing kernels (represented by circle size).

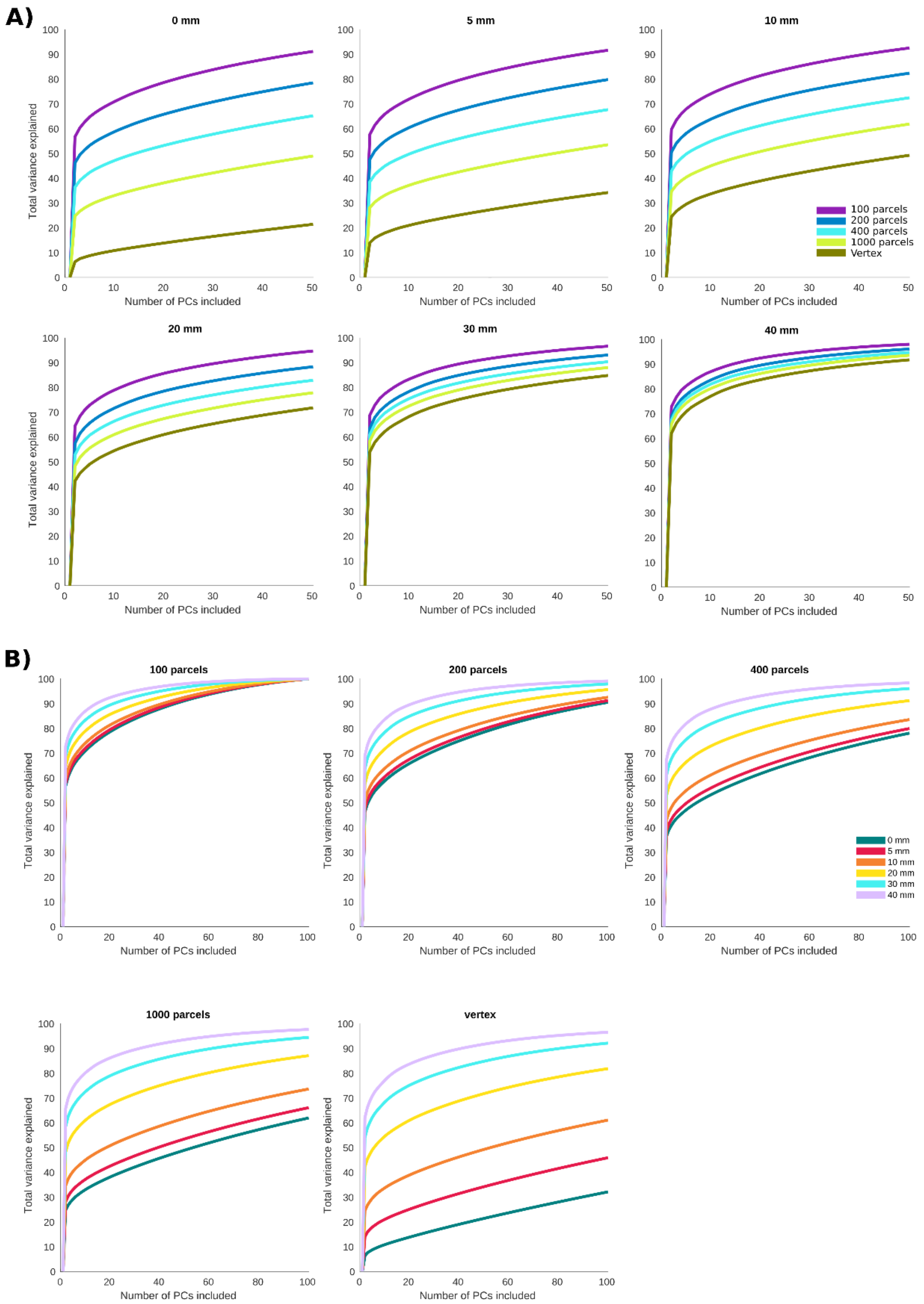


**Supplementary Figure 5.** Accumulated variance explained of cortical thickness by principal components for each smoothing/parcellation pair. **A)** results grouped together based on the smoothing level. **B)** results grouped together based on the parcellation resolution.
